# Genome diversity and quorum sensing variations in laboratory strains of *Pseudomonas aeruginosa* PAO1

**DOI:** 10.1101/2020.10.13.338434

**Authors:** Yang Liu, Stephen Dela Ahator, Yinuo Xu, Huishan Wang, Qishun Feng, Xiaofan Zhou, Lian-Hui Zhang

## Abstract

The *Pseudomonas aeruginosa* strain PAO1 has routinely been used as a laboratory model for quorum sensing (QS) studies due to its extensively coordinated regulatory circuits. However, the microevolution of *P. aeruginosa* laboratory strains resulting in genetic and phenotypic variations have caused inconsistencies in QS research. To investigate the underlying causes and impact of these variations, we analyzed 5 *Pseudomonas aeruginosa* PAO1 sublines from our laboratory using a combination of phenotypic characterization, high-throughput genome sequencing, and bioinformatic analysis. The major phenotypic variations among the sublines spanned across the levels of QS signals and virulence factors such as pyocyanin and elastase. Furthermore, the sublines exhibited distinct variations in swarming, twitching and biofilm formation. Most of the phenotypic variations were mapped to the effects of mutations in the *lasR* and *mexT*, which are key components of the QS circuit. By introducing these mutations in the subline PAO1-E, which is devoid of such mutations, we confirmed their influence on QS, virulence, motility and biofilm formation. The findings further highlight a possible divergent regulatory mechanism between the LasR and MexT in the QS pathways in *P. aeruginosa*. The results of our study reveal the effects of microevolution on the reproducibility of most research data from QS studies and further highlight *mexT* as a key component of the QS circuit of *P. aeruginosa*.

**Importance:** Microevolution of *P. aeruginosa* laboratory strains results in genotypic and phenotypic variations between strains that have a significant influence on QS research. This work highlights the variations present in *P. aeruginosa* PAO1 sublines and investigates the impact of the genetic variations on the QS circuit and QS-regulated virulence determinants. Using a combination of NGS and phenotypic analysis, we illustrate the impact of microevolution on the reproducibility of QS, virulence, motility, and biofilm studies among 5 sublines. Additionally, we revealed the significant impact of mutations in key genes such as *mexT* and *lasR* on the QS circuit and regulation of virulence. In effect, we show the need for limited propagation and proper handling of laboratory isolates to reduce the microevolution.

## Introduction

*Pseudomonas aeruginosa* causes acute and chronic infections in immune-compromised individuals and cystic fibrosis (CF) sufferers (Stover et al., 2000). Infections by *P. aeruginosa* are usually difficult to treat and persistent due to the characteristic high frequency of emergence of antimicrobial-resistant strains during therapy and the ability to switch to a biofilm state under stress conditions (Carmeli et al., 1999). As a metabolically versatile bacterium, it can adapt to myriads of environments by sensing and altering its genetic regulations to cope with imminent stress conditions. These traits have been shown to be dependent on quorum sensing (QS), the cell-density dependent regulatory mechanisms that coordinate genetic regulation in response to chemical signals or cues present in its environment (Jensen et al., 2006).

*P. aeruginosa* QS is composed of three main signals N-(3-oxododecanoyl)-L-homoserine lactone (3-oxo-C12-HSL, 3OC12HSL), N-butanoyl-L-homoserine lactone (C4-HSL, C4HSL), and 2-heptyl-3-hydroxy4(1H)-quinolone (PQS) which are produced by the *lasI, rhlI*, and *pqsABCDH* gene cluster respectively. These signals bind to their cognate regulators LasR, RhlR and PqsR(MvfR) to activate downstream virulence factors such as pyocyanin, elastase, rhamnolipids, and pyoverdine (Schuster and Greenberg, 2006; Eickhoff and Bassler, 2018). In addition, an integrative QS signal (IQS) has also been identified, which could take over the role of upstream *las* system to regulate the downstream QS systems including *rhl* and *pqs* (Lee et al., 2013). The production of signals and expression of receptors can be regulated at the transcriptional level by a series of regulators and other metabolic systems. These include the negative regulators MvaT, QscR, QslA, QteE, RpoN, RpoS, and RsaL, and positive regulators GacA/GacS, Vfr, VqsR (Lee and Zhang, 2015). The elaborate network of regulatory pathways which make up the QS circuit in *P. aeruginosa* creates a signaling continuum allowing for an effective response to varying cues which is vital for fine-tuning the adaptation of the bacterium to imminent stress conditions (Ahator & Zhang, 2019).

The adaptive processes required for the survival of *P. aeruginosa* isolates are driven by selective mutations resulting in genetic and phenotypic variations, in response to the enormous selection pressures exerted within fluctuating host environments over time (Lee and Zhang, 2015; Cordero and Polz, 2014). Spontaneous mutations in the QS systems *lasR, rhlR* and their cognate synthases *lasI* and *rhlI*, as well as other QS regulators, are frequently identified in clinical isolates (Hoffman et al., 2009). These mutations result in attenuation of virulence observed in the switch from acute to chronic infection states, biofilm to planktonic lifestyle transition, and increased fitness and growth advantage in polymicrobial settings (Wilder et al., 2011)(Köhler et al., 2009). Additionally, mutations occurring in the *gacS, retS, ampR*, and the multidrug efflux pump regulators drive the switch from acute to chronic infectious states and antimicrobial resistance (Winstanley, O’Brien, and Brockhurst, 2016; Balasubramanian, Kumari, and Mathee, 2015).

Interestingly, recent studies have also identified genetic and phenotypic diversification among the laboratory strain PAO1 from different laboratories (Klockgether et al., 2010)(Chandler et al., 2019). The studies revealed sublines of PAO1 exhibiting variability in metabolism, virulence and cell-cell signaling (Davies and Davies, 2010; Klockgether et al., 2010; Preston et al., 1995) which were proposed to arise due to prolonged propagation in selective growth media (Klockgether et al., 2010). The genetic and phenotypic diversification in the sublines have been of broad interest, due to the evidence and impact of microevolution in the sublines presented in these studies. As PAO1 is commonly used for QS research, such variations could have a significant impact on the reproducibility of research data. Although, these studies showed the phylogenetic relationship between various sublines based on their genome composition and mutations, the genetic mechanisms that influence the phenotypic variations among the sublines and their impact on the variations in QS remain vague.

To understand the underlying mechanism responsible for such variations in phenotypes of PAO1 sublines, we investigate the diversity of 5 *P. aeruginosa* PAO1 sublines and examine the effect of genomic diversity on quorum sensing through high throughput genomic sequence, virulence and bioinformatic analysis. We obtained 4 sublines from our laboratory collections and 1 subline from University of Washington (U.S.A). By applying a combination of bioinformatics and virulence assays, we were able to map the key genes (*lasR* and *mexT*) that drive the microevolution of the lab strains and also explain their potential effect on cell-to-cell signaling and virulence in *P. aeruginosa*.

## Results

### PAO1 sublines produce different levels of QS signals and QS-associated virulence factors

Phenotypic variations among the laboratory strains of *P. aeruginosa* PAO1 have been attributed to microevolution of strains during culture in selective media or prolonged passage in the laboratory(Klockgether et al., 2010)(Chandler et al., 2019). Due to the immense influence of microevolution on the repeatability of research work particularly in the field of QS, we investigated the impact of such mutations on the production of QS signals and QS associated virulence factors in 5 PAO1 sublines from our lab collection (Table 1 and S1). We examined the production of the QS signals, 3OC12HSL, PQS, and C4HSL in the PAO1 sublines using LC-MS analysis. The production of the QS signals was not consistent across the 5 sublines. The highest amount of 3OC12HSL was produced by PAO1-B and PAO1-E followed by similar levels in PAO1-C and PAO1-D (Fig 1A). On the other hand, PAO1-C and PAO1-D produced the highest level of C4HSL compared to the other three sublines (Fig 1B). No significant difference in PQS production was observed in the three sublines, PAO1-B, C, and D(Fig 1C). The subline PAO1-A was found to produce the least amount of all three QS signals, with trace amounts of 3OC12HSL detected in our analysis (Fig 1). Consistently, the relative expression levels of the QS regulators genes in PAO1-A in comparison with PAO1-E, correlated well with the production of their cognate QS signals (Fig S2).

**Figure 1.**
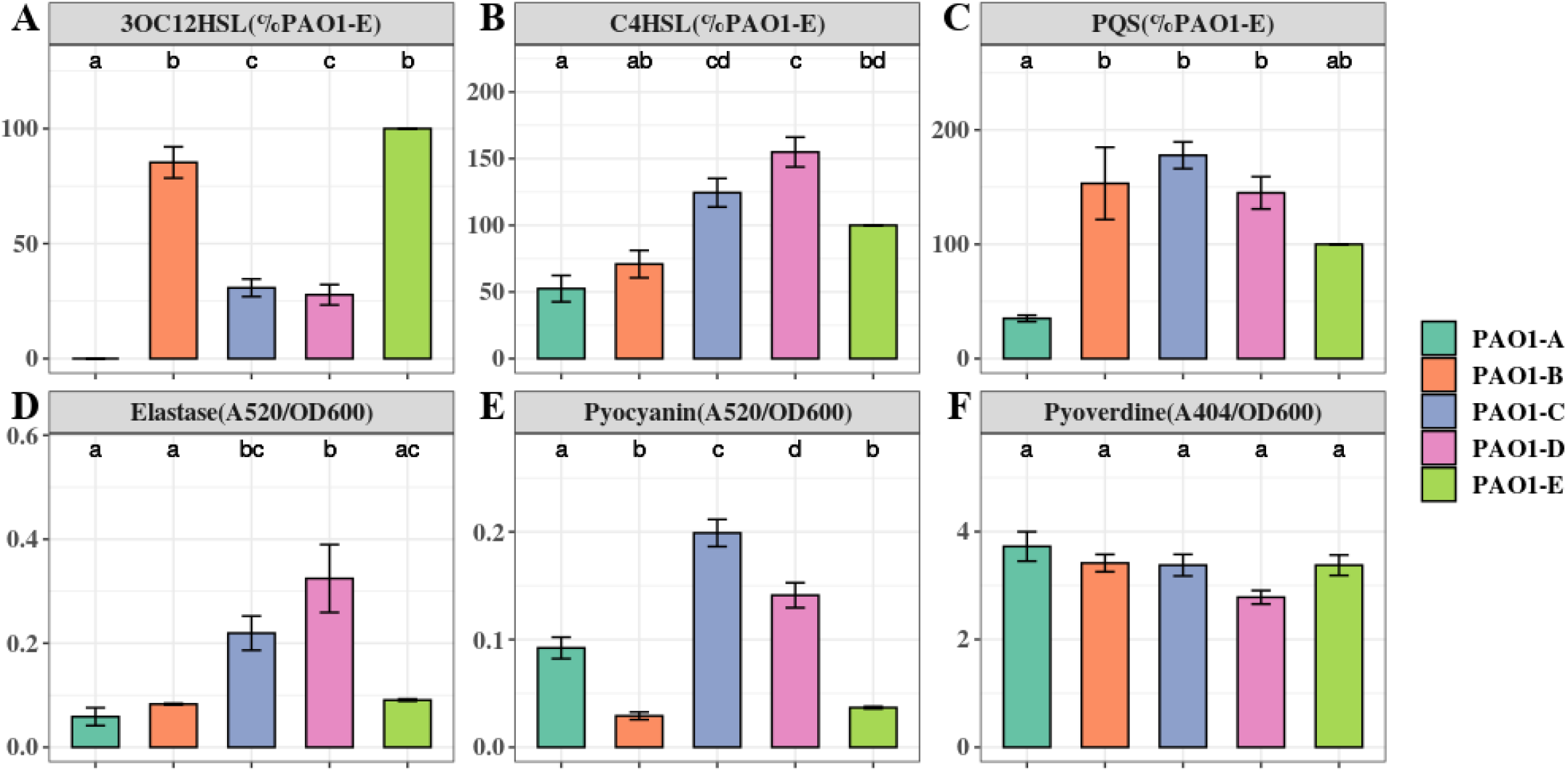
Quorum Sensing signals and virulence factors production in *P. aeruginosa* PAO1 sublines. The level of QS signals and virulence factors produced were compared to that of the PAO1-E with the mean set to 100%. Data represent the mean+/- SD of 3 independent experiments.

**Table 1.**
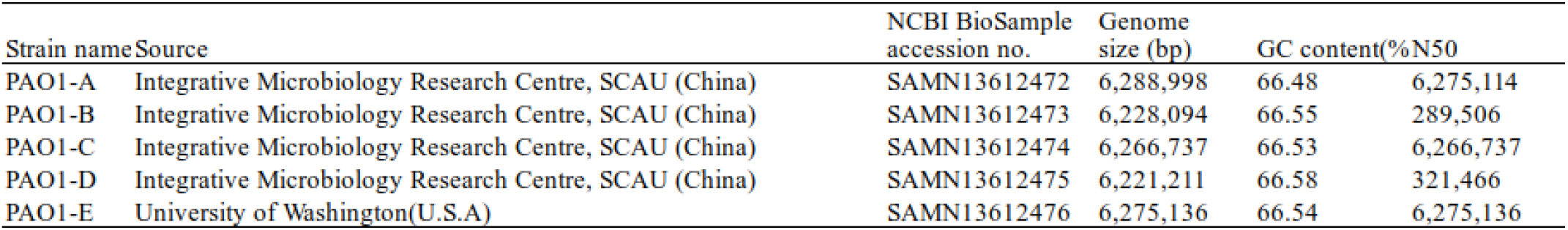
Strain selection and genome characteristics. All PAO1 subline BioSample accession numbers are listed on the table.

Elastase, encoded by *lasB* relies on the *las* QS system (Pearson, Pesci, and Iglewski, 1997). Elastase has tissue-damaging and proteinase inhibiting activity and targets plasma proteins such as immunoglobulins, coagulation, and complement factors (Pearson, Pesci, and Iglewski, 1997). Across the 5 sublines, PAO1-D produced the highest amount of elastase. However, this was not significantly greater than the amount produced by PAO1-C (Fig. 1D). Intriguingly, PAO1-A produced similar amounts of elastase as PAO1-B and PAO1-E despite its low QS signal production. (Fig 1D), Pyocyanin is an evolutionarily conserved virulence factor crucial for *P. aeruginosa* lung infection (Lau et al., 2004). Pyocyanin is regulated by the *pqs* and *rhl* QS systems, and its production is exacerbated in *lasR* mutants under phosphate depleted condition (Lau et al., 2004). Quantification of pyocyanin production revealed marked differences in levels across the 5 sublines. PAO1-C and PAO1-D produced significantly greater levels of pyocyanin compared to PAO1-B and PAO1-E (Fig 1E). PAO1-A produce pyocyanin but significantly less compared to the PAO1-C/D (Fig 1E).

Pyoverdine, the main siderophore produced by *P. aeruginosa* is regulated by QS and is a major contributor to colonization and establishing infections (Ravel and Cornelis, 2003). The result shows no significant changes in the production of the siderophore among the other 5 sublines (Fig 1F).

### Differences in biofilm formation and motility among the PAO1 sublines

Biofilm is a common adaptive state of *P. aeruginosa* which confers antibiotic resistance, enhances evasion of host immune responses, and permits persistent infections (Costerton et al., 1999; Gellatly and Hancock, 2013). Biofilm formation of the 5 PAO1 sublines was assayed using 96-well plates after a 16-hour static culture. From our biofilm assay, we observed different levels of biofilm formation across the sublines with significantly higher levels occurring in PAO1-B and PAO1-E compared with the other 3 sublines (Fig 2A).

**Figure 2.**
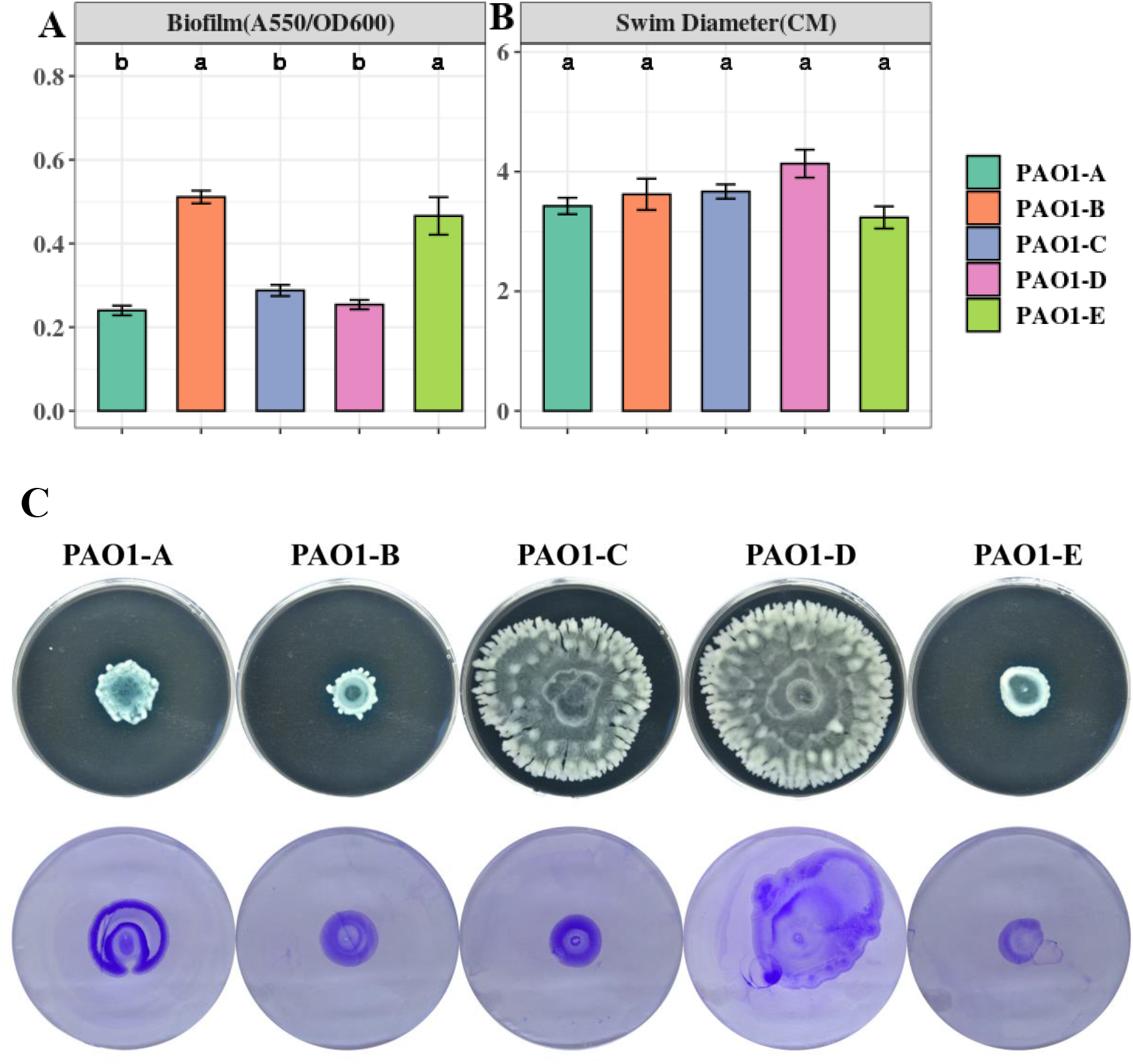
**A)** Biofilm formation was assayed from 16-hour LB cultures in 96 well plates. The data represent mean+/- SD of 6 independent biofilm assays. **B)** Swimming motility of the sublines (n=3) **C)** Swarming (top) and twitching (bottom) motility was assayed on LB medium incubated for 16 hours(n=3).

Swimming motility which is mediated by flagella was not affected by the mutations in the sublines as no significant differences were observed across the sublines (Fig 2B). However, different swarming phenotypes were observed among the strains, (Fig 2C). PAO1-D followed by PAO1-C, swarmed with the largest diameter, whereas PAO1-B and PAO1-E displayed the least swarming motility (Fig 2C) Pili formation is vital for adhesion, motility, DNA uptake, and biofilm formation (Barken et al., 2008). From our assay, pili-mediated twitching motility was inconsistent among some of the sublines. PAO1-D exhibited the highest twitching motility followed by PAO1-A. Almost similar levels of twitching were observed in PAO1-B and PAO1-C which were slightly different in comparison to PAO1-E (Fig 2C).

### Genomic variation among PAO1 sublines

To identify the underlying mutations accounting for the discrepancies in QS associated phenotypes across the 5 PAO1 sublines, we employed High-Throughput Whole Genome Sequencing (WGS) using Illunima PE-150 and Nanopore technology. The details of the whole genome sequence data of the selected *P. aeruginosa* PAO1 sublines are summarized in Table 1. The genomes of the strains were analyzed using both reference-guided mapping and *de novo* assembly approach (Fig S1), coupled with mapping and assembly-based callers for maximum variation detection.

A total of 230 SNPs and short indels was shared among the 5 sublines (Fig 3A). The detailed information of SNPs and short indels in each subline relative to the reference sequence is summarized in Table 2. Six SNPs and short indels were identified in PAO1-A genome, an 18 bp deletion in the region of *mexT*, a synonymous variant in PA0020, two resulting in missense variants of PA1975 and PA3191, one in the non-coding sequence between *psdR-dppA3* intergenic region, and one disruptive in-frame insertion of TCG in sequence for the autoinducer binding domain in LasR (Heurlier et al., 2005) (Table 2). PAO1-B and PAO1-C shared common SNPs resulting in a missense variant of the aminoacyl-tRNA biosynthesis gene, PA4277.2. Additionally, the PAO1-C subline contained the other two SNPs, one resulting in a synonymous variant in PA3316 and the other present in the non-coding region between *psdR* and *dppA3*. The latter was also identified in the PAO1-C and PAO1-D sublines (Table 2). Five other SNPs and short indels were identified in the PAO1-D genome, two synonymous variants located in *leuB* and PA0020 and another two as missense variants in *htpG* and PA3637 sequence, in addition to the intergenic region of *psdR* and *dppA3*(Table 2), also an 18 bp deletion in *mexT* that consistent with indel in PAO1-A. No unique SNPs were identified in the subline PAO1-E.

**Figure 3.**
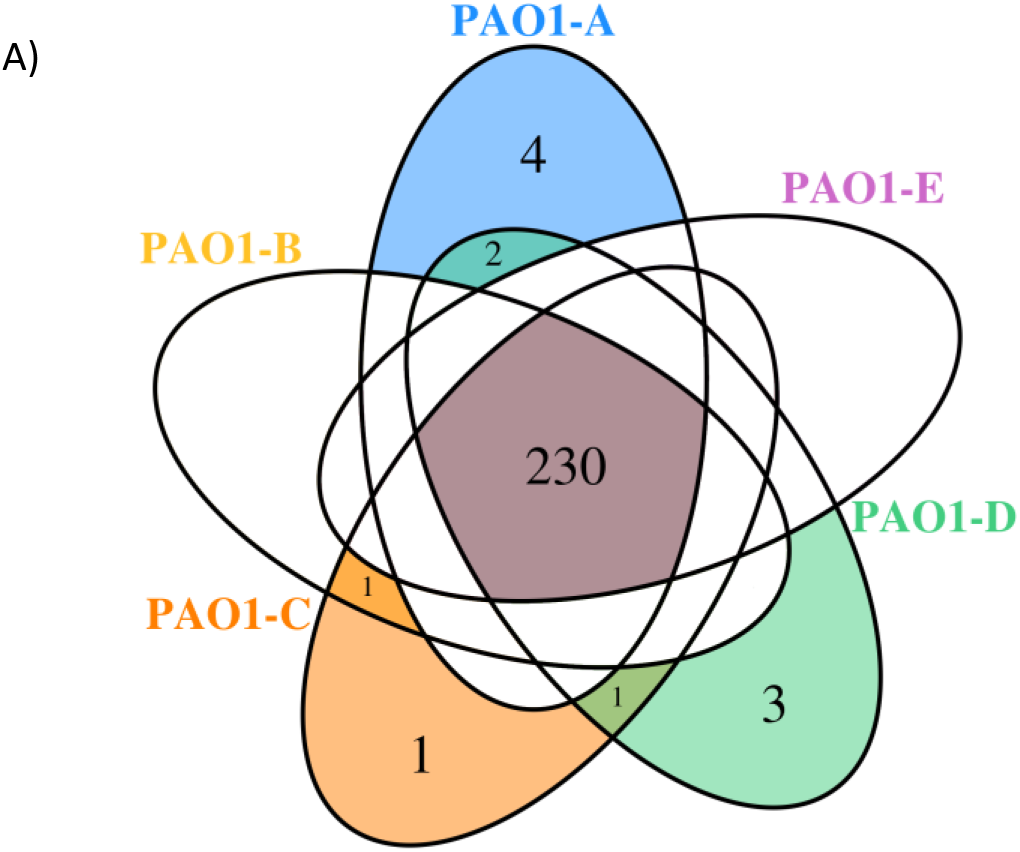
**A)** Venn diagram showing the SNPs shared among the *P. aeruginosa* PAO1 sublines.

**Table 2.**
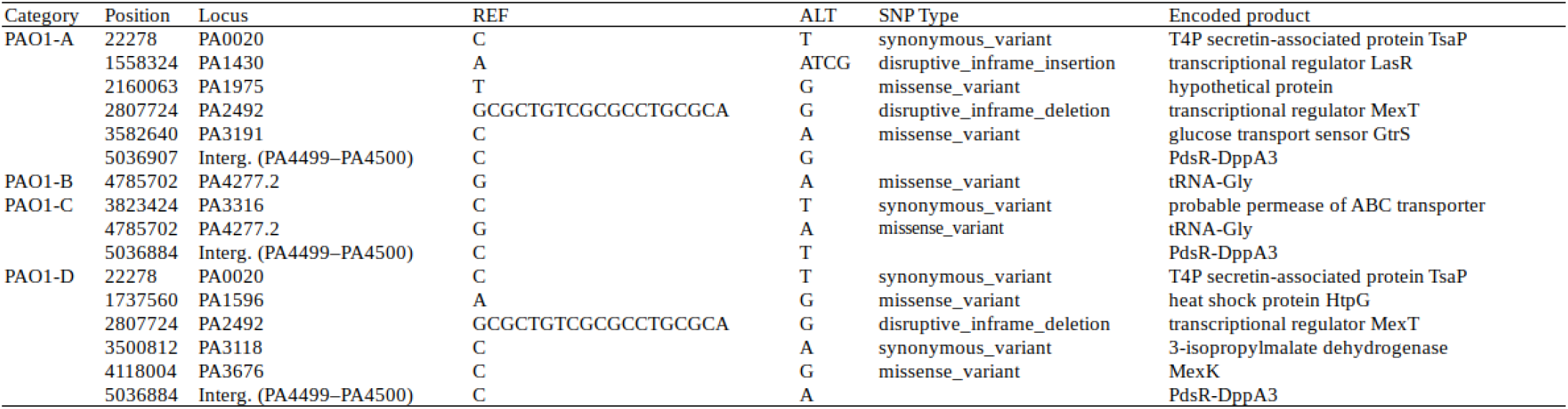
List of individual SNPs in each subline. ALT represent sublines genotype. PAO1-E is devoid of individual SNPs from whole genomic sequence.

Using comparative genomic analysis, we identified 3 structural variations (SVs), among which two were present in the 5 sublines (Table 3). These include tandem repeats or copy number variations (CNVs) occurring from PA0717-PA0727 genomic region and deletion in PA4684/PA4685 region from 5253693 to 5243687(Table 3). The PA0717-PA0727 cluster is annotated as a bacteriophage Pf1-like hypothetical protein (Hill et al., 1991), whereas PA4684, and PA4685 encode hypothetical proteins and form an operon with PA4686. These two variations in the PA0717-PA0727 genomic region and the deletion in PA4684/PA4685 region were also detected in *P. aeruginosa* PAO1-DSM (Davies and Davies, 2010), and *P. aeruginosa* isolates PA14 (Klockgether et al., 2011). One unique SV detected in PAO1-C sublines was due to 8384 bp deletion in the region containing genes of the *mexT*, Resistance-Nodulation-Cell diversion (RND) multidrug efflux (*mexE, mexF*, and *oprN*) and the downstream genes, PA2496, PA2497, and PA2498 (Table 3). This deletion in PAO1-C is located downstream of *mexT*. Also, a short indel of 18 bp was identified in both PAO1-A and PAO1-D genome in the coding sequence for the transcriptional regulator MexT (RND multidrug efflux), resulted in disruptive in-frame deletion.

**Table 3.**
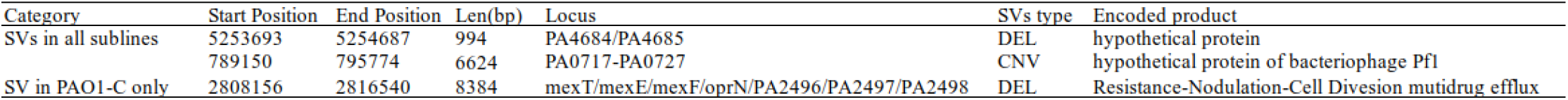
List of structure variation in each subline.

### The short Indel of *lasR* and *mexT* affect QS in *P. aeruginosa* PAO1

Mutations in *mexT* and *lasR* are commonly reported in clinical isolates and have been recently reported in lab strains and clinical isolates (Klockgether et al., 2010; Köhler et al., 2001; Kostylev et al., 2019; Sobel, Neshat, and Poole, 2005). The lasR mutants have been associated with chronic infections and increased fitness under specific metabolic stress conditions(Köhler et al., 2009). The *mexT* mutations are also known to be induced by growth in the presence of antibiotics (Sobel, Neshat, and Poole, 2005), and are associated with the regulation of most QS factors and fitness of *P. aeruginosa* strains (Kostylev et al., 2019). To further investigate the impact of the MexT and LasR on the variation of QS associated traits among the sublines, we introduced the 18bp *mexT* mutations found in PAO1-A and PAO1-D into the *mexT* of PAO1-E resulting in the strain PAO1-E4*mexT*. Additionally, we introduced the 3bp insertion in *lasR* of PAO1-A into PAO1-E to obtain the strain *PAO1-EΔlasR*. The PAO1-E4*mexT* produced a significantly decreased level of 3OC12HSL but an increased level of C4HSL production compared to the parent strain (Fig 4A, and 4B). PQS production was not significantly different between PAO1-E and PAO1-E4*mexT* (Fig 4C). Conversely, in *PAO1-EΔlasR*, the production of 3OC12HSL, C4HSL, and PQS significantly decreased in comparison with the PAO1-E(Fig 4). Additionally, in PAO1-E4*mexT* elastase and pyocyanin production levels were significantly greater than that of the parental subline PAO1-E whereas their levels significantly decreased in the *PAO1-EΔlasR* (Fig 4D, and 4E).

**Figure 4.**
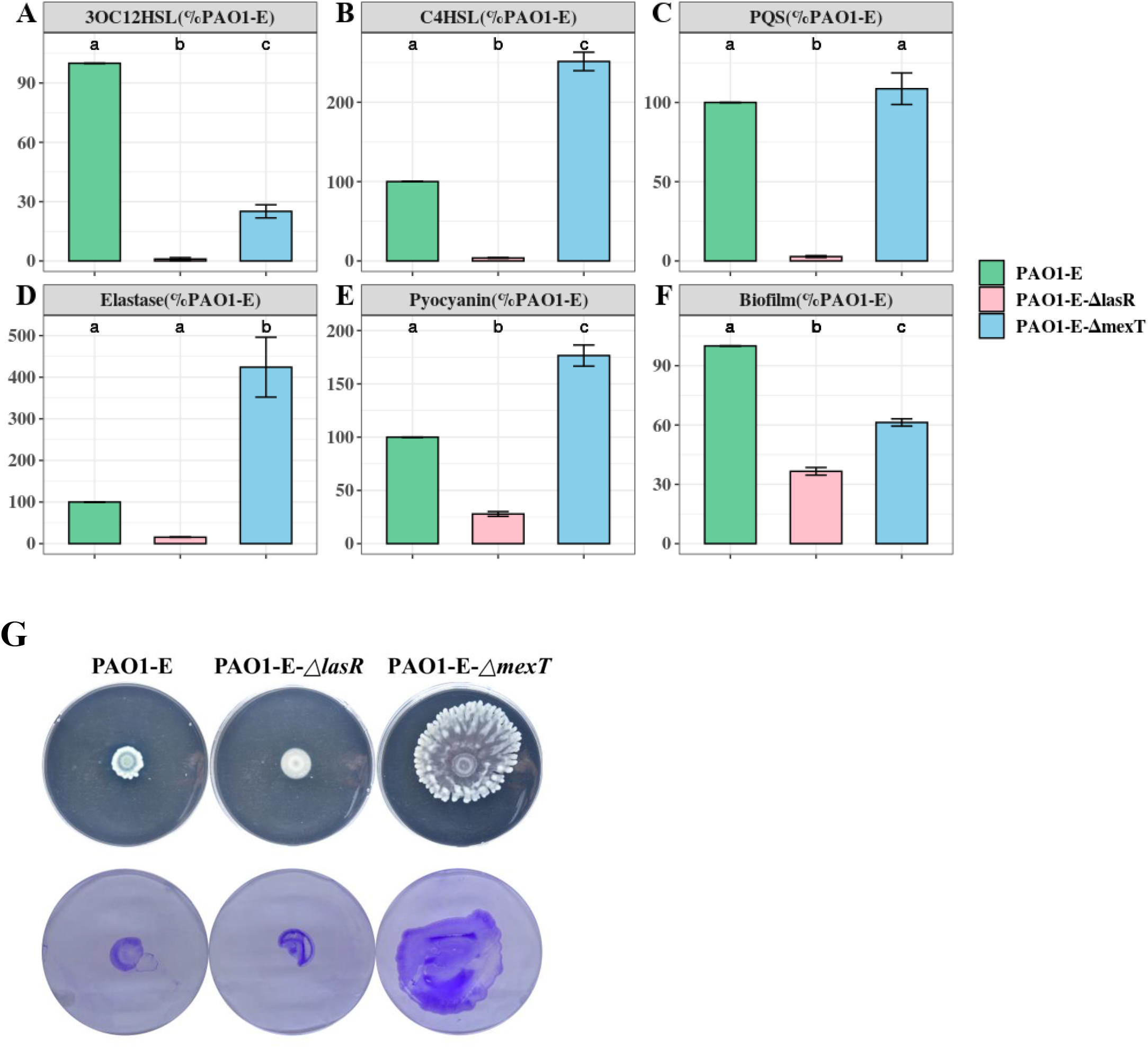
Quorum sensing signal and Virulence production in PAO1-E derivatives. **A-F)** Assay were performed with strains grown for 16 hours in LB medium. The amount of signal and virulence produced by PAO1-E was arbitrarily set at 100%. **G)** Swarming motility(top) and crystal violet stained twitching motility (bottom) of the sublines.

Both PAO1-E4*mexT* and *PAO1-EΔlasR* produced less biofilm compared to PAO1-E, however, a much significant decrease in biofilm was observed in the *lasR* mutant compared to the *mexT* mutant (Fig 4F). Although the 3bp *lasR* insertion had little effect on swarming, the 18bp deletion in *mexT* significantly increased the swarming motility in PAO1-E (Fig 4G). The introduction of the *mexT* mutations in the PAO1-E, resulted in increased twitching motility, however, no significant difference in twitching was observed in the parent strain and the *PAO1-EΔlasR* (Fig 4G).

### Evolution of *mexT* and *lasR* in *P. aeruginosa*

Based on the frequency of *lasR* and *mexT* mutations and their influence on the fitness of *P. aeruginosa* strains (Feltner et al., 2016)(Clay et al., 2020)(Kostylev et al., 2019)(Oshri et al., 2018), we decided to investigate the selective pressure driving *lasR* and *mexT* mutations in by calculating the substitution rates (nonsynonymous /synonymous (dN/dS)) of 4419 single-copy genes from 298 *P. aeruginosa* strains obtained for the Pseudomonas Genome Database(v18.1) (Winsor et al. 2015). From our analysis, we observed a higher nonsynonymous substitution rate for *lasR*, denoted by a higher dN/dS value (0.2881) in *lasR* compared to the *pqsR, rhlI, rhlR, mexT, lasI* in more than four thousand single-copy genes (Third Quartile=0.1145) (Fig 5A and Table S3). Although *mexT* is mutation-prone (Sobel et al., 2005), its low nonsynonymous substitution rates reflect a higher negative selection pressure (Fig 5A and Table S3).

**Figure 5.**
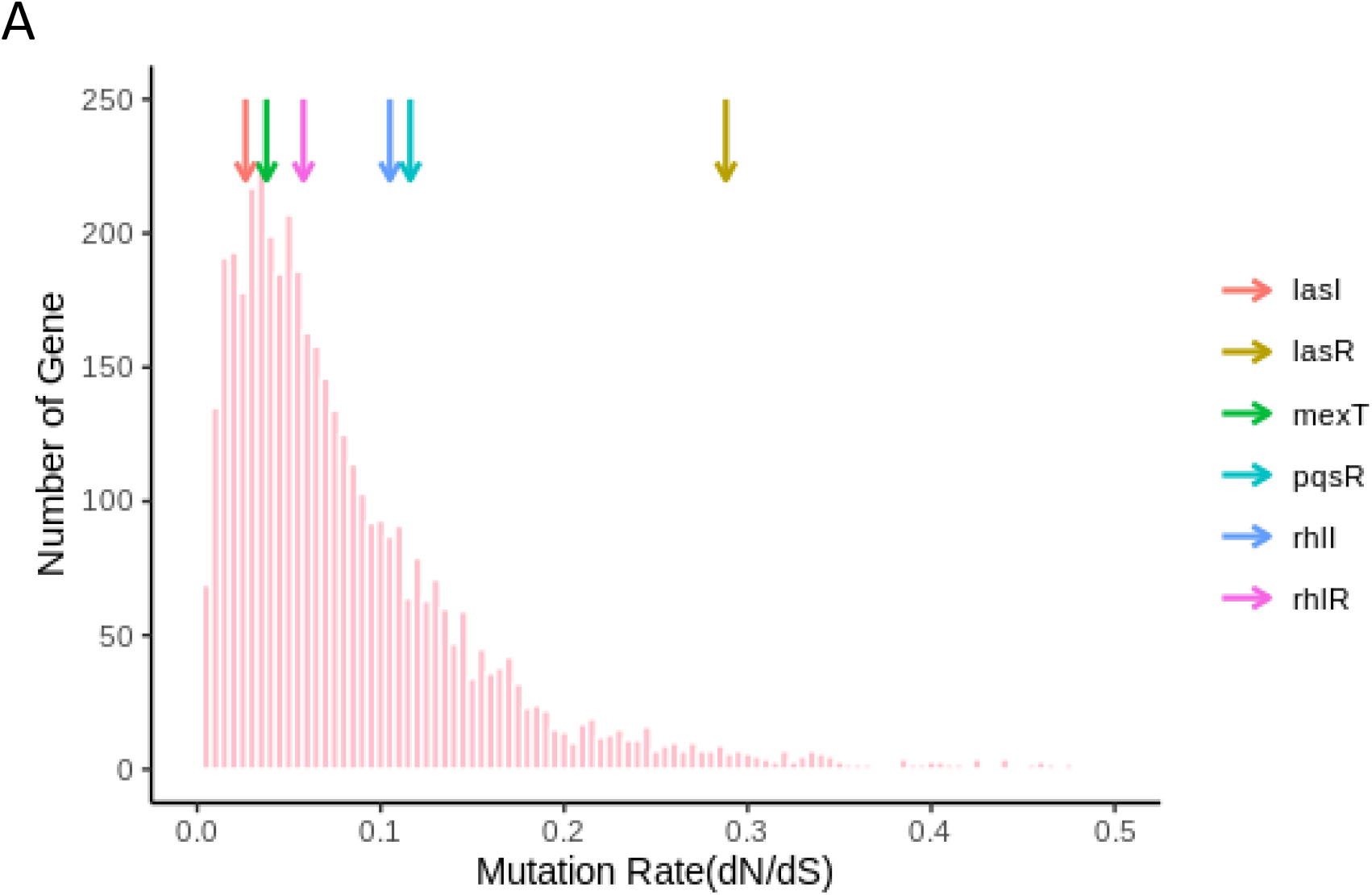
Nucleotide substitution rates of single-copy genes in *P. aeruginosa. The colored arrows show the positions of the genes and their respective dN/dS ratios*.

For further estimation of the selection pressure and the mutation hot site, we performed the codon alignment of the 2498 *lasR* sequences and 2643 *mexT* sequences and calculated the dN/dS ratio of each site (Fig 6 and Table S5, S6). Based on the mean posterior substitution rates, we observed that the LasR site shows more nonsynonymous mutation compared to the MexT in their amino acid sites (Fig 6A and Table S5, S6). In MexT, three distinct peaks at amino acid positions (17, 28, and 60) showed high mean posterior substitution rates of nonsynonymous (Fig 6A and Table S6). These results also confirmed that the *mexT* undergoes high negative selection pressure.

**Figure 6.**
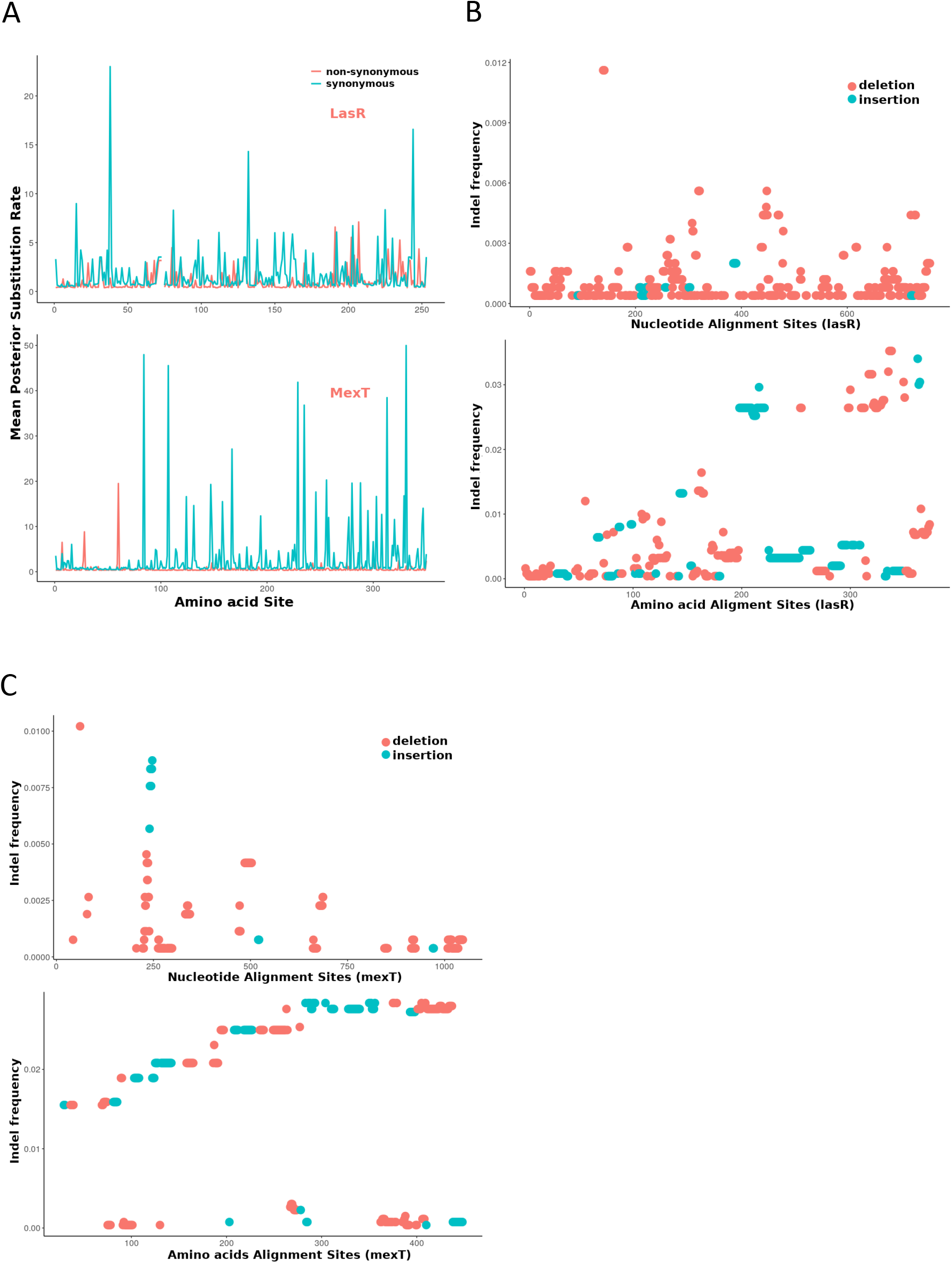
The *mexT* and *lasR* sequence substitution rate in *P. aeruginosa*. **A**. The nonsynonymous (orange peaks) and synonymous (green peaks) substitution rate in the amino acid sequences of LasR (top) and MexT (down). **B.** The indel frequency of LasR nucleotides (up) and amino acid sequences (down) in *P. aeruginosa* strains. **C.** The indel frequency of MexT nucleotides (up) and amino acid sequences (down) in *P. aeruginosa* strains. The deletion and insertion are indicated with orange and green dots.

We further investigated the nucleotide and amino acid insertion and deletion at each site of both the *lasR* and *mexT* sequences. Although the indel frequency of the *lasR* nucleotide sequences increased after 200bp with an overall higher number of deletions than insertions, the distribution of insertion and deletion was even throughout the amino acid sequences of LasR (Fig 6B).

For the *mexT* sequence, one indel-prone site(GCCGGCCAGCCGGCCA) was detected around 250bp whiles the indel frequency in the amino acid sequences of *mexT* increased from the 5’ to 3’ (Fig 6C).

## Discussion

*Pseudomonas aeruginosa* strain PAO1 is one of the most widely used model organisms for QS research. QS in *P. aeruginosa* regulates a vast majority of the physiological processes and virulence phenotypes (Ahator & Zhang, 2019) hence various research groups have focused on the development of anti-QS strategies as an alternative to combat the rising cases of antibiotic resistance in *P. aeruginosa*. However, most clinical isolates lose their QS functions via mutation in the key QS genes as well as mutation-prone genes (Hoffman et al., 2009) which makes the identification of anti-QS targets daunting. Additionally, the laboratory model organism, PAO1 from different research centers have been shown to possess gene alterations such as SNPs and deletions in some mutation hotspots which underly their phenotypic variations and influence the repeatability of *P. aeruginosa* research (Klockgether et al., 2010)(Chandler et al., 2019)(Hazen et al., 2016). Among the frequently occurring mutations in both *P. aeruginosa* clinical isolates and laboratory strains are the *lasR* and *mexT* mutations which are vital for QS regulation, multidrug resistance and drive adaptative processes in *P. aeruginosa* isolates to maximize their propagation during infection (Hazen et al., 2016)(Winstanley et al., 2016)

Although previous studies have examined the genetic and phenotypic variations arising due to microevolution in lab strains of PAO1 sublines (Klockgether et al., 2010; Chandler et al., 2019)(Hazen et al., 2016), they did not provide evidence of the underlying mutations driving the variations in QS associated phenotypes, biofilm, motility as well as other virulence determinants of the bacteria. To further understand the impact of these microevolution and the genetic basis for the variations in phenotypes among the strains in our lab, we examined the mutations present in 5 sublines of *P. aeruginosa* PAO1 and their effect on QS and virulence. Our study used a combination of whole genome sequencing and molecular biology techniques to highlights the impact of minute gene alterations on QS and virulence among *P. aeruginosa* PAO1 sublines. Significantly, we further provide evidence that mutations in the transcriptional regulators, LasR and MexT completely destabilize the QS circuit and account for the variations in the production of the PQS and C4HSL and their associated virulence factors among the sublines. Thus, indicating the significant impact of microevolution on the repeatability of QS and virulence studies using laboratory collections of PAO1.

MexT is a positive regulator of MexEF-OprN efflux pump and represses the outer membrane porin protein OprD (Sobel, Neshat, and Poole, 2005; Köhler et al., 1999; Ochs et al., 1999). From our analysis, *mexT* mutations were identified in two of the sublines with an additional subline containing deletion of the region containing the *mexT* and *mexEF-oprN* cluster as well as the PA2496, PA2497, PA2498. The *mexT* mutations have been reported in other studies of clinical and lab strains (Chandler et al., 2019; Klockgether et al., 2010; Poonsuk, Tribuddharat, and Chuanchuen, 2014; Quale et al., 2006; Sobel, Neshat, and Poole, 2005; Walsh and Amyes, 2007). In support, recent work showed MexT as a factor that reorganizes the QS system in *P. aeruginosa* independent of *lasR* and is therefore vital for the fitness of the bacteria (Kostylev et al. 2019). This in part can be due to the function of the MexEF-OprN in transporting of homoserine lactones and influence on cell-cell signaling (Köhler et al. 2001).

MexT mutations could promote pleiotropic effects on the cell as it influences the expression of at least 40 genes (Tian, Fargier, et al., 2009). Accordingly, by introducing the 18bp *mexT* mutation in the PAO1-E subline, we observed significant changes in QS signal production as well as pyocyanin, elastase, biofilm formation and motility. Based on our data, we believe that MexT may have an opposing role to LasR and may thus serve a compensatory mutation for *lasR* mutants or vice versa. PAO1-A had both *lasR* and *mexT* mutations with a characteristic loss on 3OC12HSL production but did not lose its ability to produce virulence factors such as pyocyanin, elastase and pyoverdine (Fig 4). Thus, a combination of *mexT* and *lasR* mutations does not drive the bacteria towards a non-virulent state as compared to *lasR* mutations alone. As such despite producing the least levels of QS signals with almost no 3OC12HSL, PAO1-A still produced virulence factors and formed biofilms and maintained its motility morphology comparable to the other sublines (Fig 1, 2, 4).

In support of the above observation, we note that mutation of *lasR* alone decreased the production of pyocyanin in the PAO1-E which was contrary to *mexT* mutations in the same subline. Pyocyanin is regulated in a *las*-independently manner by the *pqs* and *rhl* systems. Also, despite the defective *las* system, elastase production was comparable among PAO1-A, PAO1-B and PAO1-E. Although the *las* system regulates elastase production (Rust, Pesci, and Iglewski, 1996), the defective *las* system in PAO1-A did not cause a significant loss in elastase production. Hence it is highly possible that the defective *las* system coupled with the *mexT* mutation may account for the increase in pyocyanin and elastase production in PAO1-A. This in part could be due to the independent regulation of the *pqs* and *rhl* systems or the effect of *mexT* mutation.

Due to the importance of motility for promoting infections, colonization, and initializing biofilm formation on both biotic a nd abiotic surfaces (O’Toole and Kolter, 1998), the differences in motility observed in the sublines will greatly impact their level of pathogenicity. Another interesting observation in the interplay of *mexT* and *lasR* is the control of twitching and swarming motility. The high levels of twitching and swarming observed in the sublines, PAO1-C, PAO1-D and PAO1-E4*mexT* containing *mexT* mutations affirms the negative regulation of MexT on pili formation and flagellar mediated motility (Tian, Mac Aogain, et al., 2009). Twitching is influence by type IV pili whereas swarming is influenced by both flagellar and type IV pili (Breidenstein, Fuente-Núñez, and Hancock, 2011; Taguchi and Ichinose, 2011; Mattick, 2002; Ichinose et al., 2016). Accordingly, we believe that the increase in twitching motility in PAO1-D compared to PAO1-E is due to *mexT* mutation. Although the *las* QS system does not regulate twitching motility (Beatson et al., 2002) (Burrows, 2012), certain factors such rhamnolipids which influence motility are regulated by the *las* system (Pearson, Pesci, and Iglewski, 1997; Tian, Mac Aogain, et al., 2009; Köhler et al., 2001). The defective *lasR* and *mexT* in PAO1-A, we observe a slight increase in twitching and swarming above those of the PAO1-B and PAO1-E sublines. Also, as MexT regulation of twitching motility could be dependent or independent of MexEF-OprN (Tian, Mac Aogain, et al., 2009), we believe that the deletion of the *mexT, mexEF-oprN* gene cluster could account for the loss of twitching motility in the PAO1-C.

This work presents fascinating information about alternative pathways the compensate for loss of QS mediated functions and reaffirms the role of MexT in reorganizing the QS system in the bacteria. We observe an interesting relationship between *lasR* and *mexT*, where *mexT* tends to alleviate the loss of QS associated virulence caused by *lasR* defects which is particularly important for *P. aeruginosa* during the acute-chronic infection switch. As lower dN/dS (0.0378) in *mexT* in comparison to that of *lasR* and other single-copy genes (First Quartile=0.0351) (Fig 5A, and 6), indicated the *mexT* is under higher selection pressure. It is possible that mutations occurring in *mexT* drive the bacterial towards a more virulent state which could be compensatory and may be vital for the switch from avirulent to virulent phenotypes during the different stages of bacterial infections. Survival of *mexT* mutants is therefore cued towards the existence of synonymous mutations which favor selection or survival compared to nonsynonymous mutation.

Our study focused more on the genes that directly affect QS in *P. aeruginosa*, the interaction between *lasR* and *mexT* still need further investigation. Also mutations in genes such as *psdR* which influences the fitness of the bacteria and non-cooperative cheating in the presence of *lasR* mutants(Asfahl et al., 2015)(Kostylev et al., 2019) is currently being studies in our lab. Mutations in the intergenic region of transcriptional regulator, PsdR has been shown to arise early in the evolution of *P. aeruginosa* strains growing in the presence of casein, enhances fitness in the presence of *lasR* cheaters (Dandekar et al., 2012). DppA3 is a dipeptide binding protein with specificity for the transport of L-amino acids (Pletzer et al., 2014)(Fernández et al., 2019). Derepression of this function of DppA3 by PsdR has been shown to enhance non-cooperative cheating in *P. aeruginosa* population under QS-inducing conditions (Asfahl et al., 2015). As most of the regulatory systems in *P. aeruginosa* are highly coordinated and exhibit cross-talk, it may be a bit daunting to directly link phenotypes to specific microevolution events. Also, mutations in *lasR* and *mexT* occur during prolonged passage in special media and exposure to sub-inhibitory concentrations of antibiotics (Maseda et al., 2000; Hoffman et al., 2009), hence storage of laboratory collections of wild type PAO1 strains after prolonged passage or growth in the presence of such conditions should be avoided to minimize the microevolution of the strains. Understanding how these processes occur can help to address important problems in microbiology by explaining observed differences in phenotypes, including virulence and resistance to antibiotics and the discrepancies in QS research.

## Materials and Methods

### Bacterial strains and growth conditions

The *P. aeruginosa* PAO1 strains used in this study are list in Table 1. All Strain were maintained in 40 % glycerol, 60 % Lysogeny Broth (LB, 1 /L, 15 g Agar, 10 g Tryptone (Sigma-Aldrich), 5 g Yeast Extract(Sigma-Aldrich), 10 g NaCl) at −80 °C. For all experiments, cultures were inoculated directly from the stock used for sequencing without subculturing.

### Genomic DNA extraction and whole genomic sequences

The EasyPure Bacteria Genomic DNA Kit (EE161-01, Transgenbiotech, Beijing, China) was used for the extraction of the genomic DNA from the sublines. The concentration of genomic DNA was measured by NanoDrop and stored at −20 °C. The genomic DNA of the five sublines were submitted for sequencing using the Illumina NovaSeq S4 PE-150 (Novogene, China) and Oxford Nanopore MinlON (Nextomics Biosciences, China).

### Genome assembly, mapping and genome annotation

Sequences were checked by FastQC software (Andrews et al., 2010), a quality control tool for high throughput raw data. Short reads were mapped against the reference using Burrows-Wheeler Aligner BWA-MEM (Li and Durbin, 2009) whereas long reads were mapped using Minimap2 (Li, 2018). *De novo* assembly was performed using Unicycler (Wick et al., 2017) with SPAdes algorithm and assembled data summarized by BBMap (Bushnell, 2014). For gene prediction and annotation, the DDBJ Fast Annotation and Submission Tool DFAST (Tanizawa, Fujisawa, and Nakamura, 2017). pyani(Pritchard et al., 2016) software ANIb method were used to phylogenetic analysis by calculating Average Nucleotide Identity (ANI).

### SNP detection and analysis

A combination of software was used for SNPs calling. The SAMtools and bcftools (Li, 2011) were used to map short reads aligned with the *P. aeruginosa* PAO1 reference genome (NO. NC_002516.2). GATK Best Practices (Van der Auwera et al., 2013; DePristo et al., 2011) was used for variant calling workflow. The SAMtools and bcftools calling were trimmed by removing MIN(QUAL) < 100 SNPs. GATK SNPs calling were followed by germline short variant discovery (SNPs + Indels) using HaplotypeCaller and GenotyperGVCFs tools (Poplin et al., 2018). SNPs were further trimmed by removing MIN(QUAL) <500. SNPs were also identified in the assembled data generated by the Unicycler using Mummer (Kurtz et al., 2004) and progressiveMauve (Darling, Mau, and Perna, 2010). The SNPs produced by the four tools were merged and false-positive SNPs eliminated by checking the original mapped short reads bam file manually using IGV (Robinson et al., 2011). The supporting reads which were less than 25 % were not considered as SNPs.

### Structure variation (SV) detection and analysis

Integrated structural variant multiple callers were used to detecting SVs. The Structural Variants from Mummer svmu (Chakraborty et al., 2018) tool was used to compare *de novo* assembly sequence against the reference. BreakDancer (Chen et al., 2009) was used to set sorted mapping input bam files and filter the total number of reads pairs > 3 or confidence score > 85 %. Using Pindel (Ye et al., 2009), which operates on a read-pair based method, the outputs allele depth(AD) over 20 % were kept and for split-reads based DELLY (Rausch et al., 2012), the outputs paired-end supported site(PE) < 2 were discarded. Also, Svseq2 (Zhang, Wang, and Wu, 2012) was used to detect deletions and insertions. All results were merged to obtain a final list of SVs by a union of the output from the individual callers. A diagrammatic representation of the filter parameter is shown in Fig S1.

### SNPs and SV annotation

The common SNPs and SVs in the sublines were manipulated by the command line script to separate from individual variation. All SNPs and SVs were customized into a VCF file on demand by the shell script. SnpEff (Cingolani et al., 2012) was used for variation annotation to predict the effect of the generic variants against the SnpEff database *P. aeruginosa* PAO1 strain.

### Selective pressure of *Pseudomonas aeruginosa* single-copy genes

The raw data (Pseudomonas Ortholog Groups) used for the mutation rate analysis was collected from Pseudomonas Genome Database (Winsor et al., 2015)(v18.1). The downloaded Ortholog files were filtered by the python script to obtain only 4419 single-copy gene and 298 Pseudomonas aeruginosa strain. The nucleotide sequences were extracted by mapping gene name and strain name to Pseudomonas Genome Database (Winsor et al., 2015)(v18.1) Annotations (GFF3) and Genomic DNA (Fasta) files. Single-copy gene files were translated by EMBOSS Transeq tool (Rice, Longden, and Bleasby, 2000), aligned by mafft tool (parameter:retree 1) (Katoh et al., 2002). The codon alignment was generated through pal2nal.pl (Suyama, Torrents, and Bork, 2006) program and the alignment files were trimmed by trimAl (parameter:gappyout) (Capella-Gutiérrez, Silla-Martínez, and Gabaldón, 2009). The required treefile for subsequent analysis was generated by iqtree (parameter:st=DNA m=GTR+G4 nt=1 fast) (Nguyen et al., 2014) for single-copy gene files, individually. The mutation rate of each single-copy gene was calculated by HyPhy-Branch-Site Unrestricted Statistical Test for Episodic Diversification (hyphy BUSTED) (Murrell et al., 2015). Nonsynonymous/synonymous (dN/dS) ratio were generated by improving branch lengths, nucleotide substitution biases, and global dN/dS ratios under a full codon model. The mutation rates are in Table S3.

### Estimate mean posterior synonymous substitution rate and mutational type of *P. aeruginosa lasR* and *mexT* gene site

The *lasR* and *mexT* sequences were blasted against all *P. aeruginosa* complete and draft genome in the Pseudomonas Genome Database (Winsor et al. 2015)(v18.1). Codon alignment of the *lasR* (2498) and *mexT* (2643) sequences was performed by transeq, mafft and pal2nal tools. The multiple sequence alignments files were trimmed by the python script to make them inframe and remove the stop codon. The mutation rate of each site was calculated by hyphy FUBAR(Murrell et al. 2013). The mutational types were calculated via a Biopython script.

### In-frame deletion and knock-in

DNA manipulation was conducted by In-frame deletions and insertion described previously (Filloux and Ramos, 2014). The DNA fragments for 3bp insertion in *lasR* and 18bp deletion in *mexT* mutations were synthesized by Sangon Biotech (China). The fragments were cloned into pK18mobsacB plasmid using ClonExpress MultiS One Step Cloning Kit (C113-01, Vazyme) for construction gene knock-in and deletion constructs. The constructs were transformed into *E. coli* S17-1 for conjugation with PAO1-E. Transconjugants were selected on Minimal Media(MM) supplemented with gentamicin (30 μg/mL) and transferred onto MM supplemented with 10% (wt/vol) sucrose to select mutants. Mutants containing the desired deletion and insertion were confirmed by PCR and DNA Sanger sequencing.

### Motility

Motility was assayed by Plate-Based method as previously described (Filloux and Ramos, 2014). Swimming motility was assessed on 0.3 % agar plates (1 /L, 3 g Bacto agar (Becton Dickinson), 8 g Nutrient Broth (Becton Dickinson)). Overnight cultures (37 °C, 200 rpm; LB) were used to inoculate swim plates by depositing 1 μl of culture directly into the agar in the center of the plate. Plates were incubated face up at 37 °C, and the swim diameter (in centimeters) recorded at 16h. Swamming motility was assessed on 0.6 % agar plates (1 /L, 6 g Bacto agar (Becton Dickinson), 5 g Bacton-peptone (Becton Dickinson), 3 g Yeast Extract (Sigma), 5 g D. glucose). Overnight cultures (37 °C, 200 rpm; LB) were used to inoculate swam plates by depositing 1 μl of culture directly into the agar in the center of the plate. Plates were incubated face up at 37 °C, and the swam recorded at 16h. Twitching motility was assessed on 1.5 % agar LB plates. Overnight cultures (37 °C, 200 rpm; LB) were used to inoculate twitch plates by depositing 1 μl of culture directly into agar in the bottom of the plate. Plates were incubated face down at 37 °C for 16h, and the Twitch visualized by fixing the culture with Water: Glacial acetic Acid: Methanol at a ratio of 4: 1: 5 and stained with 0.1% crystal violet.

### Pyoverdine quantification

*Pseudomonas aeruginosa* PAO1 were cultivated in 37 °C in Iron-depleted succinate medium(1 /L, 7.86 g K_2_HPO_4_·3H_2_O, 3 g KH_2_PO_4_, 1 g (NH_4_)_2_SO_4_; 0.1 g MgSO_4_·7H_2_O; 4 g succinate; PH=7.0) (Stintzi et al., 1998). The OD600 was recorded after 24 h culture using spectrophotometer. Cell-free supernatant was collected by max speed centrifuged and measured at A404 was recorded using succinate medium as a blank.

### Pyocyanin quantification

Pyocyanin was assayed from *P. aeruginosa* PAO1 cultured in LB medium overnight at 37 °C and 250 rpm. Single colony was inoculated into 10 mL culture for 16 h. The 5 mL culture were centrifuged at 12,000 × *g* for 5 min and the cell free supernatants mixed with equal volume of chloroform followed by continuous rocking for 30 min at room temperature. The solvent phase was obtained by brief centrifugation, mixed with 5 mL 0.2 mol/ L HCl and rocked at room temperature for an additional 30 min (Filloux and Ramos, 2014). The pyocyanin quantification was determined by measuring absorbance of supernatant at A520 nm and normalizing against the cell density at OD600.

### Elastase quantification

Elastase production in *P. aeruginosa* strains were performed by Elastin-Congo Red (Sigma) assay (Ohman, Cryz, and Iglewski, 1980). Single colonies of the *P. aeruginosa* strains were inoculated into 10 mL LB and cultured for 16 h at 37 °C and 250 rpm. The cultures were centrifuged at 12,000 × *g* for 5 min to obtain cell-free supernatant. Briefly, 500 μL of bacterial cell-free supernatant was mixed with an equal volume of 5 mg/ mL elastin-Congo red with ECR buffer in 2 mL Eppendorf tube and incubated at 37 °C shaker for 2 h. The quantity of Congo red dye released from the elastin digestion is proportional to the amount of elastase in the supernatant. Elastase quantification was determined using a spectrophotometer at A520 and normalized against the cell density at OD600.

### Biofilm formation assay and quantification

Biofilm formation was assayed by 96-well plates as previously described (Filloux and Ramos, 2014). A single colony was inoculated into 10 mL LB broth and grown at 37 °C, 200 rpm overnight. OD600 was measured by nanodrop spectrophotometer and the culture was diluted to OD600 = 0.5. A volume of 1 μL diluted cells was added to 200 μL LB medium in sterile 96 well plate incubate at 37 °C statically for 16 h. The plate was washed with Ultra-pure water at least 3 times and stained with 250 μL 0.1 % crystal violet for 15 min. The plate was rinsed, dried at room temperature and the remaining dye was solubilized with 300 μL Dimethyl sulfoxide (DMSO). The dissolved biofilm was measured by the spectrophotometer at absorbance A550.

### Quorum sensing signal extraction and quantification

QS signal extraction was conducted as previously described by (Dong et al., 2008). Single colonies of the *P. aeruginosa* cells were inoculated into 5 mL LB broth and grown overnight at 37 °C and 200 rpm. The signals were extracted from 5 mL of supernatants with an equal volume of acidified ethyl acetate (0.1 % Acetic acid) twice. The organic phase was transfer to a fresh tube and dried with nitrogen gas. The extracted compounds were dissolved in 1 mL filtered HPLC grade methanol for LC-MS analysis.

The LC-MS method was adapted from the Nishaben M. Patel method (Patel et al., 2016). HPLC was performed on a Dionex UltiMate 3000 system (Thermo Fisher Scientific) using a C18 reverse-phase column (Thermo Fisher Scientific) and varying concentration gradients of methanol and consisted of 0.1 % acidified water as mobile phase. The gradient profile for chromatography was as follows: 2 % methanol and 98 % water for 1.5 min, linear increase in methanol to 100 % over 5 min, isocratic 100 % methanol for 4 min, and then equilibration with 2 % methanol and 98 % water for 1.5 min. The flow rate was constant at 0.4 mL/ min.

Compounds separated by HPLC were detected by heated electrospray ionization coupled to high-resolution mass spectroscopy (HESI-MS, Q Exactive Focus, Thermo Fisher Scientific). The analysis was performed under positive ionization mode. Settings for the ion source were: 10 aux gas flow rate, 40 sheath gas flow rate, 0 sweep gas flow rate, 4 kV spray voltage, 320 °C capillary temperature, 350 °C heater temperature, and 50 S-lens RF level. Nitrogen was used as a nebulizing gas by the ion trap source. The MS/MS method was designed to perform an MS1 full-scan (100 to 1000 m/z, no fragmentation) together with the SIM model. Settings for the SIM method were 35000 resolution, 1.0 m/z isolation offset, 4.0 isolation window and centroid spectrum. Signals mass scans were set 3OC12HSL at 298.20128 m/z, C4HSL at 172.09682 m/z, PQS at 260.1645 m/z, respectively. Data analysis was performed using the Thermo Xcalibur software (Thermo Fisher Scientific) and TraceFinder (Thermo Fisher Scientific).

### RNA purification and qPCR analysis

Overnight culture of *P. aeruginosa* PAO1 were diluted in LB broth and incubated at 37 °C to OD600 =1.0. Bacterial pellets were obtained by centrifugation at 4 °C for 3 min at 12,000 × *g*. Total RNA samples were purified using the RNeasy miniprep kit (Z3741, Promega) following the manufacturers’ instruction. Genomic DNA was digested by using the TURBO DNA-free Kit (AM1907, Thermo Fisher Scientific) and the integrity and purity of the RNA determined by nanodrop and gel electrophoresis. cDNA was generated by using FastKing RT Kit (KR116, Tiangen, China) and Real-time qPCR was carried out using PowerUp™ SYBR™ Green Master Mix (A25742, Applied Biosystems™) in the QuantStudio™ 6 Flex Real-Time PCR System (Applied Biosystems™). The *proC* and *rpoD* were used as house-keeping genes. The primer specific to the original copy of genes are list in the table S2.

### Data analysis

Data are expressed as means ± standard error. Significance was determined using one-way ANOVA analysis of variance with Tukey HSD multiple comparisons in Python(version 3.7). A P value of < 0.05 was considered significant. Plots were generated by R (version 3.60).

### Accession number(s)

This Whole Genome Sequence project has been deposited at NCBI/DDBJ/ENA under the BioProject accession number PRJNA596099.

## Supplemental Material

**M 1. Supplemental Table.**

**M 2. Supplemental Figure.**

## Acknowledgments

This work was supported by the Natural Research Foundation of China (Grant No.: 31330002), Key Projects of Guangzhou Science and Technology Plan (Grant No.: 201804020066), Guangdong Technological Innovation Strategy of Special Funds (Grant No.: 2018B020205003).

## Notes

### Competing Interest Statement

The authors have declared no competing interest.

