## Supplemental Table and Figure for "Genome diversity and quorum sensing variations in laboratory strains of *Pseudomonas aeruginosa* PAO1": 20200310_lyon_Supp_Figures.pptx

### Slide 1
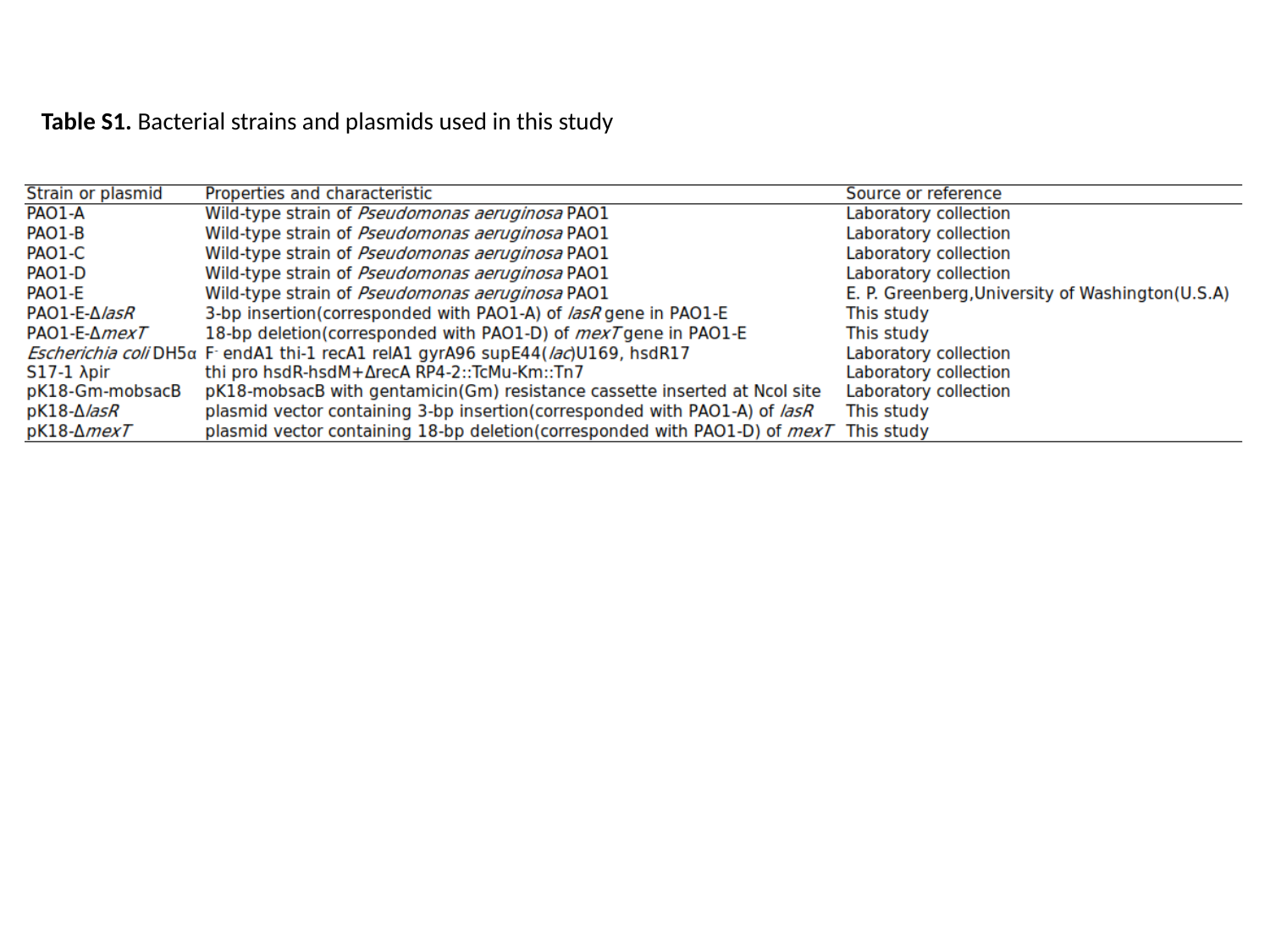

Table S1. Bacterial strains and plasmids used in this study

### Slide 2
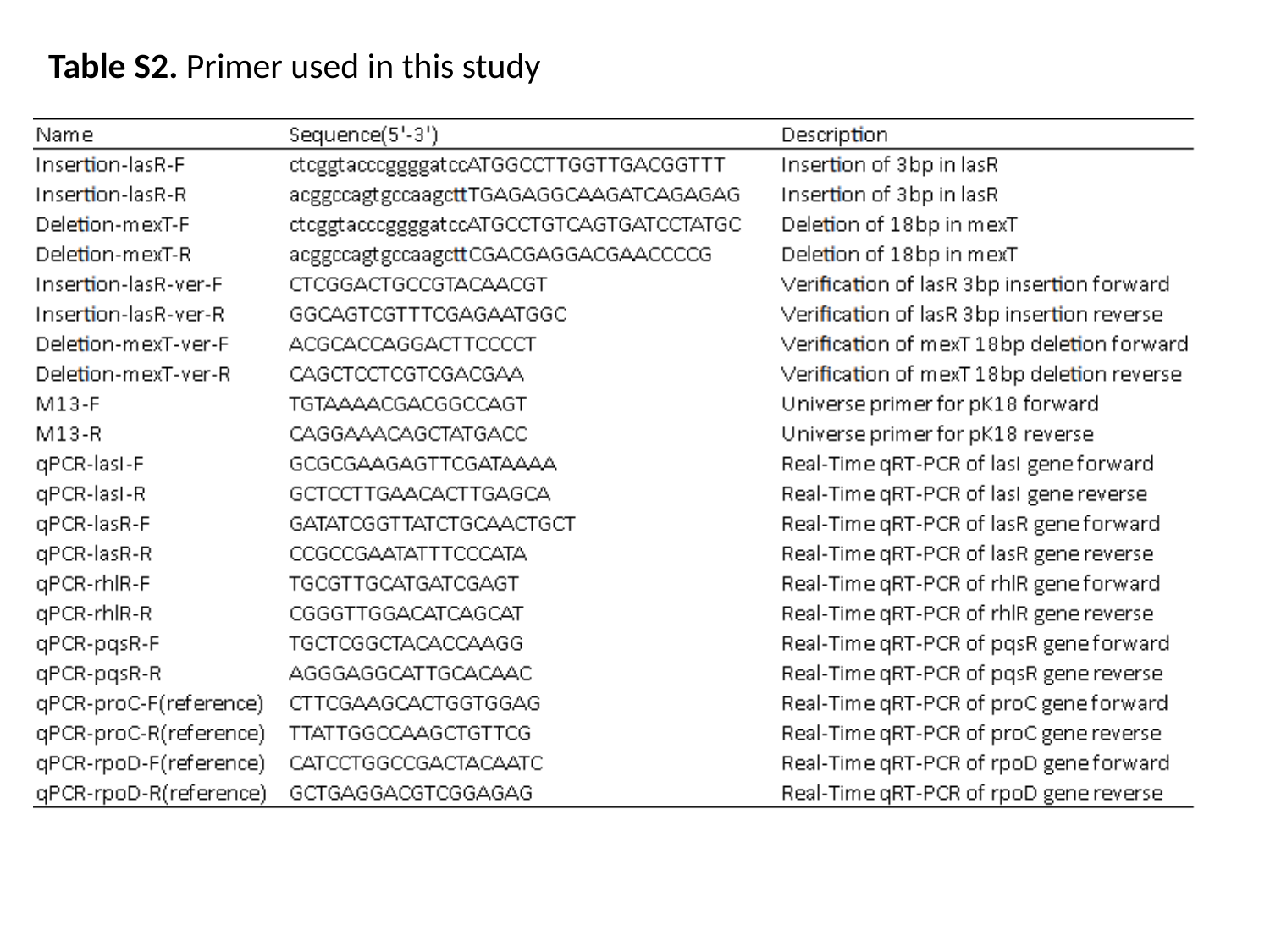

Table S2. Primer used in this study

### Slide 3
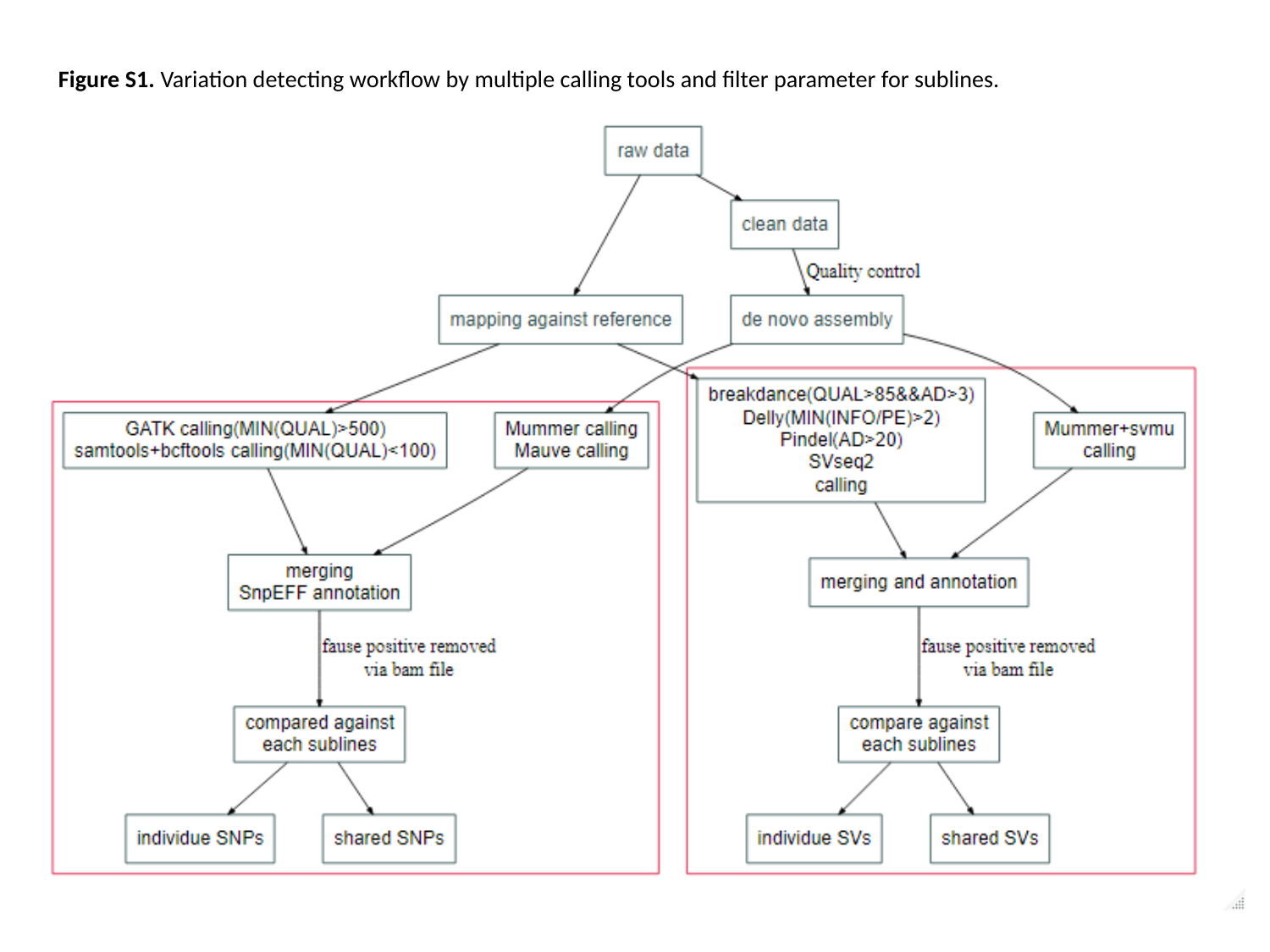

Figure S1. Variation detecting workflow by multiple calling tools and filter parameter for sublines.

### Slide 4
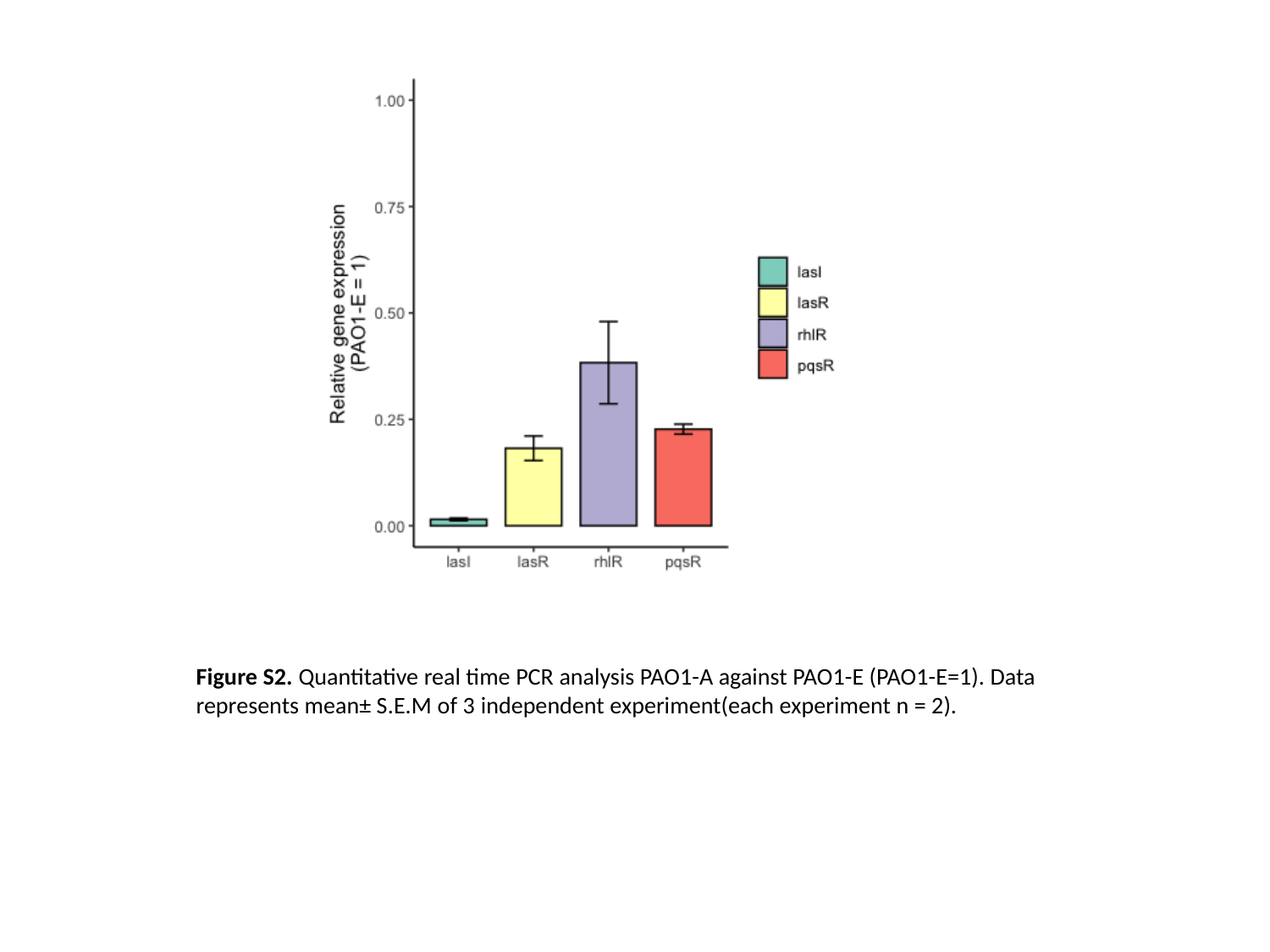

Figure S2. Quantitative real time PCR analysis PAO1-A against PAO1-E (PAO1-E=1). Data represents mean± S.E.M of 3 independent experiment(each experiment n = 2).
